# Differences in cognitive functions between cytomegalovirus-infected and cytomegalovirus-free university students: a case control study

**DOI:** 10.1101/275792

**Authors:** Veronika Chvátalová, Blanka Šebánková, Hana Hrbáčková, Petr Tureček, Jaroslav Flegr

## Abstract

Cytomegalovirus (CMV) is the herpetic virus, which infects 45 – 100% people worldwide. Many reports suggest that CMV could impair cognitive functions of infected subjects. Here we searched for indices of effects of CMV on infected subjects’ intelligence and knowledge. The Intelligence Structure Test I-S-T 2000 R was used to compare IQ of 148 CMV-infected and 135 CMV-free university students. Infected students expressed higher intelligence. Paradoxically, their IQ decreased with decreasing concentration of anti-CMV antibodies, which can be used, statistically, as a proxy of the time passed from the moment of infection in young subjects when the age of subjects is statistically controlled. The paradox of seemingly higher intelligence of CMV infected subjects could be explained by the presence of the subpopulation of about 5 – 10 % CMV-positive individuals in the population of “CMV-negative students”. These false negative subjects had probably not only the oldest infections and therefore the lowest concentration of anamnestic antibodies, but also the lowest intelligence among the infected students. Prevalence of CMV infection in all countries is very high, approaching sometimes 90 %. Therefore, the total impact of CMV on human intelligence may be large.

## Introduction

Cytomegalovirus (CMV) is a ubiquitous virus belonging to the *Herpesviridae* family with high prevalence rates; 45 – 100 % women of reproductive age are infected worldwide ^1^. In immunocompetent patients, this infection is assumed as asymptomatic; it, however, causes dramatic complications in immunocompromised subjects. Congenital cytomegalovirus infection is considered the main infectious cause of brain damage, cognitive delay and sensorineural hearing loss worldwide ^2^. Following primary infection, CMV establishes lifelong latent infection with possible reactivation and reinfection. Latent phase in immunocompetent individuals is usually considered as asymptomatic. However, infection with CMV has been associated with immunosenescence, functional impairment, frailty, cardiovascular disease and Alzheimer’s disease ^3-8^. Postnatal acquired CMV infection has also been associated with impaired cognition, predominantly in specific populations, i.e., in elderly with cardiovascular disease ^9^, healthy elderly ^10,11^, schizophrenics ^12,13^ and AIDS patients ^14^. The association of asymptomatic CMV infection with cognitive performance in healthy individuals has come into research focus only recently ^15-17^.

The main aim of the present study was to search for indices of impaired cognitive functions in university students with anamnestic anti-CMV antibodies. We measured the intelligence in a cohort of 283 biology students of the Faculty of Science, Charles University. In the double blind study, we searched for possible differences in intelligence between CMV-infected and CMV-free students and for possible correlations between intelligence and concentration of anti-CMV IgG antibodies using standard IQ test, The Intelligence Structure Test I-S-T 2000 R.

## Materials and methods

### Subjects

All undergraduate students enrolled for courses of Evolutionary biology and Practical Methodology of Science in 2010 – 2012 were invited by e-mail to participate voluntarily in the research projects studying effects of parasites on human behaviour, performance and personality. About 60 – 70 % of invited students signed an informed consent form and provided 5 ml of blood (taken by medical personnel) for serological analysis. Testing of intelligence proceeded about 6 months after recruitment during the years 2010 and 2012. In time of IQ testing, neither the probands, nor researchers were aware about CMV status of particular subjects. The study was conducted in accordance with the Declaration of Helsinki, and the protocol was approved by the IRB of the Faculty of Science, Charles University (2013/07). All subjects were adult and all signed a written informed consent form approved by the IRB.

### Intelligence testing

Testing of intelligence was performed with the Czech version of The Intelligence Structure Test I-S-T 2000 R ^18,19^. The test consists of a basic module, which comprises of three verbal, three numerical, and three abstract figural reasoning subtests and a knowledge test. The knowledge test is focused on questions from geography / history, business, arts/culture, mathematics, science and daily life. Using both the basic module and the test of knowledge enabled us to obtain a broad spectrum of results: verbal (VI), numerical (NI) and figural intelligence (FigI); verbal (VK), numerical (NK), figural (FigK) and general knowledge (GK); fluid (FI), crystallized (CI) and general intelligence (GI). The test was administered on computers to groups of 7 – 12 individuals in the same room and at the same time (9:15 am). The total length of the test was about 145 minutes including a 15 minutes break before administering the second module with the knowledge test.

### Immunological tests for CMV

A sandwich ELISA method with inactivated CMV AD 169 strain antigen (ETI-CYTOK-G PLUS, DiaSorin, Saluggia, Italy) was used for determination of CMV status of students (CMV-infected vs CMV-free). This method enables quantitative detection CMV IgG from cut-off value 0.4 IU / ml to 10 IU / ml and expresses excelent specificity and very good sensitivity ^20^. All subjects with ambiguous results of the serological test, i.e., with IgG concentration 0.36 – 0.44 IU / ml were excluded from the analyses.

### Statistical analysis

The SPSS 21^®^ program was used for statistical testing (frequency tables, general linear models) and to check statistical tests assumptions. Nonparametric tests were calculated with partial Kendall test ^21,22^; using an Excel sheet available at http://web.natur.cuni.cz/flegr/programy.php (item 12).

In the descriptive statistics, including in the figure, we used age standardized IQ scores. However, in all statistical tests we used raw IQ scores (sums of the correct answers) as the dependent variables and controlled for the effect of age (and sex and the age-sex interaction in the parametric tests) by including these confounding variables as predictors in our statistical models. This is the only possible approach when specific populations (e.g. students, people of a specific narrow age strata, etc.), and not the general population, are being studied. Even if population norms for a specific population under study are available (which is not true for biology students), it is always better to use raw data (and control for age and sex statistically) in cross-sectional studies to avoid potential problems with cohort effects and to also solve the problem of continuously changing norms in our rapidly changing social and technical environment.

A partial Kendall correlation test allows one to control for only one variable. Therefore, we included the age of a subject as a parameter functioning as the continuous covariate, and we controlled for sex by performing separate analyses for male and female students. In the parametric tests – MANCOVA and ANCOVA general linear model (GLM) tests – with the components of intelligence as the dependent (outcome) variables (continuous) and CMV seropositivity status (binary variable) or the concentration of anti-CMV IgG antibodies (continuous variable) as the focal predictor, and sex (binary), age (continuous), and the sex-age interaction as the confounding variables. As the MANCOVA test with all components of intelligence showed a significant association between intelligence and CMV infection, we did not use any correction for multiple tests in the follow-up ANCOVA tests. The correlation between IQ components and age were studied with general linear models (ANCOVA) and only the results significant after the correction for multiple tests by Benjamini-Hochberg procedure were printed in bold in the table. The false discovery rate in permutation tests (pre-set to 0.2) was controlled with Benjamini-Hochberg procedure ^23^. In contrast to the simple Bonferroni’s correction, this procedure also takes into account the distribution of p values of performed multiple tests.

### Permutation tests

Permutation tests were performed using the program TREEPT (PTPT) ^24,25^ modified for an analysis of data contaminated with unknown number of subjects with false negative diagnosis using the method of reassignment of potentially false negative subjects ^26^. This freeware program is also available at http://web.natur.cuni.cz/flegr/treept.php. The algorithm of the permutation test with data reassignment was as follows: Particular percentage (5, 10, 15, 20 or 25 %) of subjects with the lowest values of the dependent variables – here the standard scores of particular components of intelligence were relocated from the group of 135 CMV-negative subjects to the second group of 148 CMV-positive subjects. Then, the difference of means of these two groups was calculated. In the next 9,999 steps, the empirical values of the analyzed variable were arbitrarily assigned into two groups of 135 and 148 cases, the particular percentage of cases with the lowest values of the variable (intelligence) in the smaller group were relocated to the larger group, and the difference of the means of the two groups was calculated. Finally, all 10,000 differences (including the one calculated from non-permuted data) were sorted from highest to lowest. The percentage of the differences higher or equal to that calculated on the basis of non-permuted data was considered as the statistical significance (p), i.e., the probability of obtaining the same or higher difference of means of groups of 135 and 148, if subjects were assigned into these groups randomly before the relocation of subjects with lowest intelligence from the smaller to the larger group. The results of a Monte Carlo simulation using R 3.3.1 for Windows show that the permutation test for contaminated data could not provide false positive results, i.e., could not provide a more significant result than a standard permutation test if no false negative subjects exist in the population under study, see the Results, part f.

### Availability of materials and data

The datasets generated during and/or analysed during the current study are available in the figshare repository, https://figshare.com/s/97b37051afb7545819b8.

## Results

### a) Descriptive statistics

Two hundred eighty-three students of the Faculty of Science, Charles University, 197 women (21.2 years, S.D. = 1.9) and 86 men (21.5 years, S.D. = 2.0) were tested for specific immunity against CMV. The prevalence rates of CMV infection was 52.2 % for all, 49.7 % in women and 58.1 % in men (Chi^2^ = 1.69, p = 0.194). The prevalence of CMV infection fluctuated depending on size of settlements where subjects spent their childhood from 47 % to 59 %. However, these differences were not significant (all: Chi^2^ = 2.25, p = 0.689, women: Chi^2^ = 4.87, p = 0.300, men: Chi^2^ = 3.05, p = 0.548). Similarly, no relation was observed between intelligence and the size of settlements where subjects spent their childhood (all: F_1,467_ = 0.010, p = 0.919, women: F_1,292_ = 0.472, p = 0.493, men: F_1,172_ = 0.187, p = 0.174). Average IQs for each component of intelligence are shown in Table 1. We detected significant negative (women) and positive (men) associations between age and some components of intelligence (Table 2).

**Table 1.** Means and standard deviations of all components of intelligence in standard scores for women and men analyzed together and women and men analyzed separately.

|  | Total sample<br>(N=283) |  | Women<br>(N=197) |  | Men<br>(N=86) |  |
| --- | --- | --- | --- | --- | --- | --- |
|  | Mean | Std. Dev. | Mean | Std. Dev. | Mean | Std. Dev. |
| Verbal intelligence | 131.32 | 7.85 | 130.89 | 7.21 | 132.30 | 9.12 |
| Numerical intelligence | 110.88 | 12.23 | 109.36 | 11.40 | 114.36 | 13.37 |
| Figural intelligence | 111.21 | 13.22 | 110.95 | 13.49 | 111.80 | 12.62 |
| Verbal knowledge | 119.20 | 9.65 | 117.76 | 9.35 | 122.51 | 9.55 |
| Numerical knowledge | 120.99 | 12.05 | 119.02 | 11.30 | 125.51 | 12.56 |
| Figural knowledge | 120.34 | 10.04 | 118.69 | 10.20 | 124.13 | 8.61 |
| General knowledge | 122.73 | 9.37 | 120.84 | 8.89 | 127.05 | 9.05 |
| Crystallized intelligence | 133.12 | 9.70 | 131.25 | 9.23 | 137.40 | 9.44 |
| Fluid intelligence | 123.65 | 12.47 | 122.77 | 11.95 | 125.65 | 13.45 |
| General intelligence | 120.16 | 10.00 | 119.20 | 9.59 | 122.35 | 10.62 |

**Table 2.** Correlations between IQs and age.

|  | Total sample (N=283) |  |  | Women (N=197) |  |  | Men (N=86) |  |  |
| --- | --- | --- | --- | --- | --- | --- | --- | --- | --- |
| | p | $\eta^2$ | B | p | $\eta^2$ | B | p | $\eta^2$ | B |
| Verbal intelligence | 0.453 | 0.002 | 0.104 | 0.837 | 0.000 | 0.032 | 0.370 | 0.010 | 0.254 |
| Numerical intelligence | 0.241 | 0.005 | -0.341 | **0.027** | 0.025 | -0.739 | 0.385 | 0.009 | 0.495 |
| Figural intelligence | 0.100 | 0.010 | -0.383 | **0.032** | 0.023 | -0.617 | 0.783 | 0.001 | 0.109 |
| Verbal knowledge | 0.092 | 0.010 | 0.147 | 0.396 | 0.004 | 0.090 | 0.086 | 0.035 | 0.267 |
| Numerical knowledge | 0.428 | 0.002 | -0.078 | **0.006** | 0.038 | -0.316 | 0.022 | 0.061 | 0.422 |
| Figural knowledge | 0.729 | 0.000 | -0.030 | 0.501 | 0.002 | -0.074 | 0.646 | 0.003 | 0.063 |
| General knowledge | 0.854 | 0.000 | 0.039 | 0.240 | 0.007 | -0.301 | 0.044 | 0.047 | 0.752 |
| Crystallized intelligence | 0.658 | 0.001 | 0.678 | 0.460 | 0.003 | -1.365 | 0.068 | 0.039 | 4.963 |
| Fluid intelligence | 0.094 | 0.010 | -3.097 | **0.008** | 0.036 | -5.711 | 0.499 | 0.005 | 2.387 |
| General intelligence | 0.212 | 0.006 | -0.620 | **0.023** | 0.026 | -1.324 | 0.362 | 0.010 | 0.858 |

Values of p, η^2^, B (the slope of a correlation line, positive for the positive relations between the cognitive performance and age) refer to age as computed with GLM. The resuls significant after the correction for multiple tests are printed in bold.

### b) Effect of CMV on intelligence – multiple multivariate test

Multiple multivariate GLM analyses, with independent variables CMV and age (and also sex, sex-CMV and sex-age interactions when the total sample was analysed) and verbal, numerical, figural intelligence, verbal, numerical, figural knowledge as dependent variables, performed separately for the total sample, women and men showed a significant positive association of CMV and intelligence in the total sample (p<0.001; η ^2^ = 0.085) and in women (p = 0.010; η ^2^ = 0.084). This association was not significant in men (p = 0.057; η ^2^ = 0.142). Besides that, an association between sex and intelligence (p = 0.047; η ^2^ = 0.045) and between the interaction sex-age and intelligence (p = 0.023; η ^2^ = 0.052) was observed in the total sample; in women there was a significant negative correlation between age and intelligence (p = 0.007; η ^2^ = 0.089).

### c) Effects of CMV on particular components of intelligence – post hoc tests

Simple multivariate GLM analyses were performed with particular components of intelligence (VI, NI, FigI, VK, NK, FigK, GK, CI, FI and GI, respectively – see Table 3) as dependent variables and CMV and age (and also sex, sex-CMV and sex-age interactions when the total sample was analysed) as independent variables. Because of the positive result of previous multiple multivariate test, no correction for multiple statistical tests was performed. These post hoc tests revealed significantly higher IQ scores in verbal intelligence (p = 0.039; η ^2^ = 0.015) and verbal knowledge (p<0.001; η ^2^ = 0.049) in CMV-infected subjects compared to CMV-free subjects in the total sample. The results also showed that women scored worse than men on numerical knowledge (p = 0.002; η ^2^ = 0.035). In the total sample, the age was positively associated with verbal knowledge scores (p = 0.045; η ^2^ = 0.014); interaction sex-age was associated with numerical knowledge (p<0.001; η ^2^ = 0.044), general knowledge (p = 0.024; η ^2^ = 0.018), fluid intelligence (p = 0.043; η ^2^ = 0.015) and general intelligence (p = 0.046; η ^2^ = 0.014). Visual inspection of data showed that these scores decreased with the age in women and increased with the age in men.

**Table 3.** Intelligence of CMV infected and CMV-free subjects.

|  | Total sample<br>(N=283) |  | Women<br>(N=197) |  | Men<br>(N=86) |  |
| --- | --- | --- | --- | --- | --- | --- |
|  | CMV-free | CMV-infected | CMV-free | CMV-infected | CMV-free | CMV-infected |
| Verbal intelligence | 130<br>(8.4) | 132<br>(7.2) | 130<br>(7.0) | 131<br>(7.4) | 132<br>(11.6) | 134<br>(6.6) |
| Numerical intelligence | 110<br>(12.0) | 112<br>(12.4) | 108<br>(11.0) | 114<br>(11.8) | 110<br>(13.6) | 115<br>(13.3) |
| Figural intelligence | 111<br>(12.7) | 111<br>(13.7) | 111<br>(12.6) | 113<br>(14.4) | 111<br>(13.1) | 111<br>(12.4) |
| Verbal knowledge | 117<br>(9.2) | 121<br>(12.1) | 116<br>(9.0) | 119<br>(9.4) | 120<br>(9.6) | 125<br>(8.9) |
| Numerical knowledge | 121<br>(12.0) | 121<br>(12.1) | 119<br>(11.1) | 124<br>(11.6) | 119<br>(13.9) | 127<br>(11.5) |
| Figural knowledge | 121<br>(10.1) | 120<br>(10.1) | 119<br>(10.0) | 124<br>(10.6) | 118<br>(9.7) | 124<br>(7.8) |
| General knowledge | 122<br>(9.2) | 124<br>(9.4) | 120<br>(8.6) | 125<br>(9.2) | 121<br>(10.1) | 128<br>(8.1) |
| Crystallized intelligence | 132<br>(9.6) | 134<br>(9.7) | 131<br>(9.2) | 135<br>(9.3) | 132<br>(10.2) | 139<br>(8.7) |
| Fluid intelligence | 123<br>(12.3) | 124<br>(12.6) | 122<br>(11.6) | 126<br>(12.3) | 124<br>(13.8) | 126<br>(13.3) |
| General intelligence | 119<br>(9.7) | 121<br>(10.2) | 118<br>(9.1) | 122<br>(10.1) | 120<br>(11.0) | 123<br>(10.4) |

The table shows means and standard deviations (in parentheses).

When analyzed separately for women and men, CMV-infected women scored significantly higher than CMV-free women in verbal knowledge (p = 0.005; η ^2^ = 0.040) and CMV-infected men obtained higher scores than controls in verbal knowledge (p = 0.009; η ^2^ = 0.080).

Certain components of intelligence were not normally distributed even after transformations (Shapiro-Wilks test of Log transformed variables: verbal intelligence: p<0.001; verbal knowledge: p = 0.009). Therefore, we also tested the influence of CMV on intelligence with partial Kendall test, the nonparametric test allowing controlling for one confounding variable. Models included CMV and age as independent variables and particular components of intelligence (VI, NI, FigI, VK, NK, FigK, GK, CI, FI, GI, respectively) as dependent variables. The results obtained using these models were similar to those obtained using GLM analysis. There were significant positive associations between CMV and verbal intelligence (p = 0.035; τ = 0.084), verbal knowledge (p<0.0005; τ = 0.195), crystallized intelligence (p = 0.020; τ = 0.093), and general knowledge (p = 0.018; τ = 0.094) in the total sample. When analyzed separately for women and men, CMV was positively associated with verbal knowledge in women (p = 0.001; τ = 0.160) and in men (p = 0.001; τ = 0.240). CMV-infected subjects obtained higher scores on these components of intelligence compared to uninfected controls.

### d) Correlation between intelligence and concentration of anti-CMV antibodies

The level of specific IgG antibodies fluctuates in time depending on physiological status of an individual and on environmental factors. However, it generally declines with time from infection even in pathogens with dormant stages in nervous tissue ^27^. Positive relation between age and concentration of anti-CMV antibodies as well as elevated level of these antibodies before deaths of CMV-infected seniors have been described in literature ^28,29^. However, no relation (in women) or even the negative relation (in men) between age and concentration of specific anti-CMV antibodies was shown (but was not discussed) in CMV-infected subjects of age 1 – 20 (women) and 1-24 (men) years – graph 2 in ^28^. Additionally, no relation was shown between the concentration of antibodies and time since the infection in a longitudinal study performed on 18 women of reproduction age ^30^ and a longitudinal study performed on 20 plasma donors demonstrated that eleven donors vacillated at least once between seronegative and seropositive statuses during a period of 16 months, more frequently to the seronegative status (40 vs 24). The authors showed in detail the course of fluctuations (with generally negative trends) of anti-CMV antibodies in three donors and concluded that a negative serology cannot be equated with a lack of prior host contact with CMV ^31^. Therefore, it can be assumed (statistically) that the university students infected a long time ago would have on average lower levels of specific antibodies than the subjects infected recently. If changes in intelligence are caused by the infection, differences in IQ scores between CMV-infected and CMV-free subjects should gradually increase with decline of antibodies level. Correlations of the level of specific IgG anti-CMV antibodies with each component of intelligence were analyzed with partial Kendall test. Models included concentration of specific IgG anti-CMV antibodies and age as independent variables and dependent variables always were one of the ten components of intelligence (VI, NI, FigI, VK, NK, FigK, GK, CI, FI, GI). Positive correlations between the level of IgG antibodies and scores obtained on verbal (p = 0.044; τ = 0.112) and fluid intelligence (p = 0.034; τ = 0.118) were observed in the CMV-infected subsample when men and women were analyzed together. No significant correlations were observed in the CMV-infected subsample when analyzed separately for men and women, see Fig. 1 and Fig. 2. In the CMV-free subsample there was significant negative correlation between the level of IgG antibodies and IQs on numerical knowledge (p = 0.042; τ =-0.118) when men and women were analyzed together. When analyzed separately for men and women, negative correlation between the level of IgG antibodies and IQs on figural intelligence was observed only in men (p = 0.047; τ =-0.232). We did not search for correlations between the detected concentration of anti-CMV antibodies in the whole, CMV-unsorted population as two different phenomena, positive correlation of IQ with the concentration of specific anti-CMV IgG antibodies and negative correlation of IQ with the concentration of cross-reacting antibodies of an unknown specificity were observed in CMV seropositive and CMV seronegative subpopulations, respectively.

**Figure 1.**
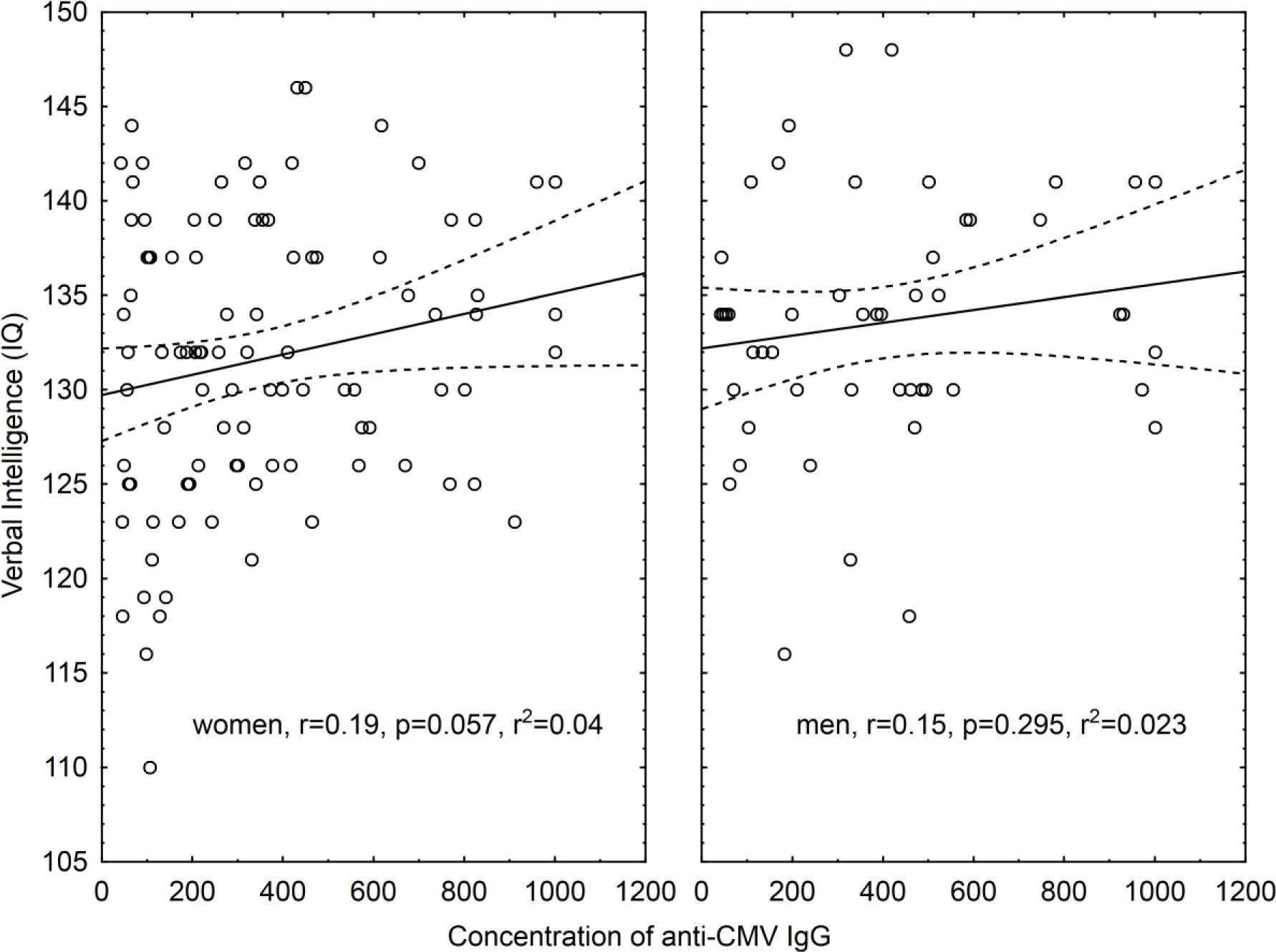
Correlation between antibody titre and verbal intelligence in CMV-positive subjects. The abscissa shows verbal intelligence of women and men in standard scores and the ordinate shows the level of specific antibodies in arbitray units deffined by the manufacturer of the ELISA kit.

**Figure 2.**
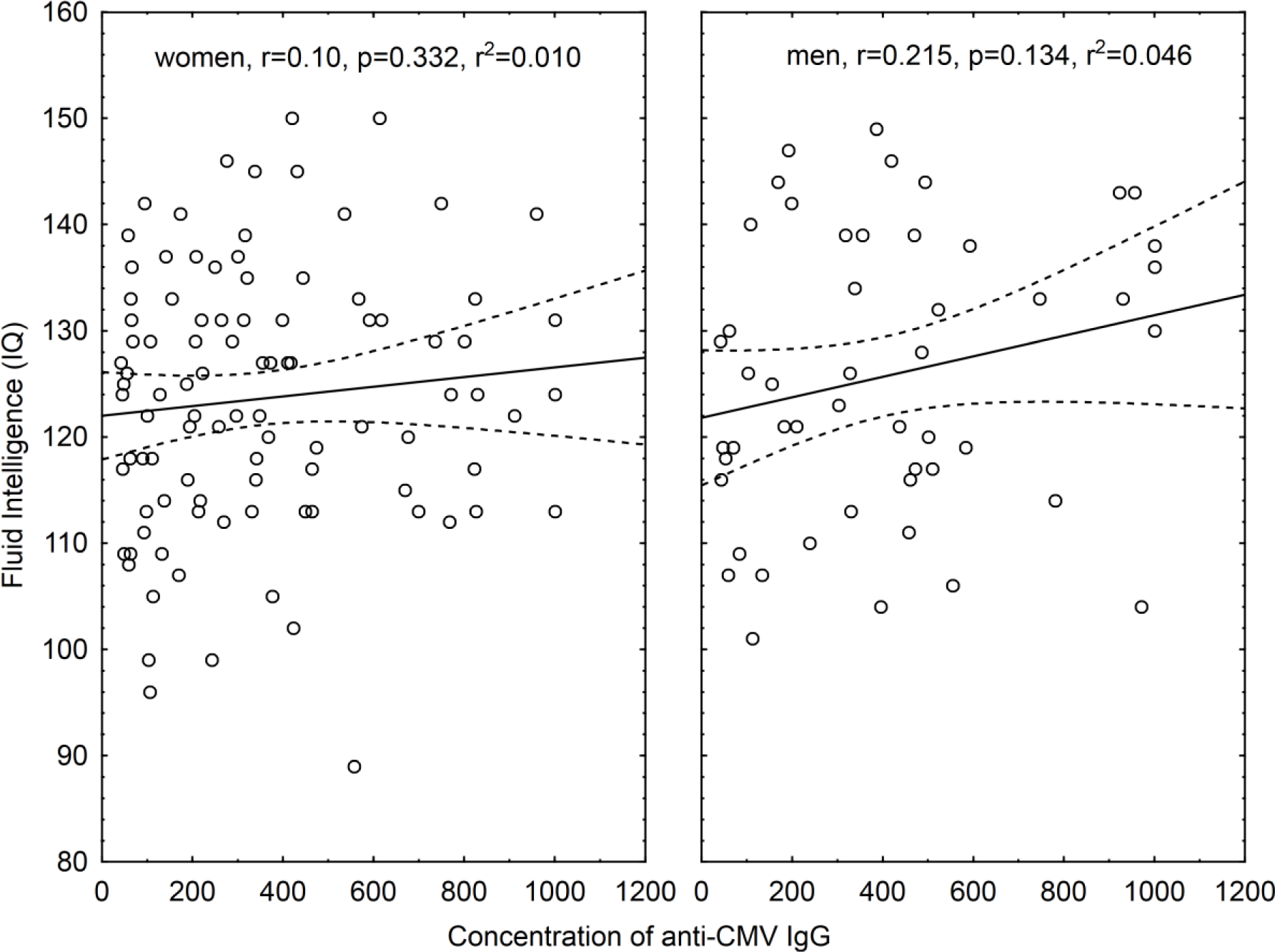
Correlation between antibody titre and fluid intelligence in CMV-positive subjects.

The abscissa shows fluid intelligence of women and men in standard scores and the ordinate shows the level of specific antibodies in arbitray units deffined by the manufacturer of the ELISA kit.

### e) Differences in IQ between CMV-positive and CMV-negative subjects estimated by permutation tail probability test with data reassignment

The reported decrease of specific antibodies with time from infection increases the risk of false negative test results in students with an old CMV infection, e.g., in individuals infected in early childhood. Our subsample of CMV-free subjects could be therefore contaminated with an unknown number of misdiagnosed CMV-positive individuals that have been infected a long time ago ^26,32^. This subpopulation of CMV-infected but CMV seronegative subjects could be the most influenced by CMV and could have the lowest IQ scores because of their long duration of infection or because of their infection in early stages of ontogenesis, see the Fig. 3. That could explain the observed paradox, i.e., infected subjects have on average higher IQ scores and at the same time the intelligence of infected subjects declines with the length of infection assessed by the level of IgG antibodies. To check the premises of present model we examined the IgG titres of 24 CMV-seropositive subjects who have been tested by us two times within an interval 5-83 months. In 8 subjects the concentration of specific anti-CMV IgG antibodies increased while in 16 subjects decreased. The nonparametric Wilcoxon matched pairs test showed that this decrease was significant (T = 72, Z = 2.23, p = 0.026). One of these 24 subjects turned seronegative between the first and the second test. Contamination of CMV-free subsample with false negative individuals can be revealed and its impacts on results of statistical tests eliminated by permutation tests with reassignment of suspect cases between subsamples ^26,32^.

**Figure 3.**
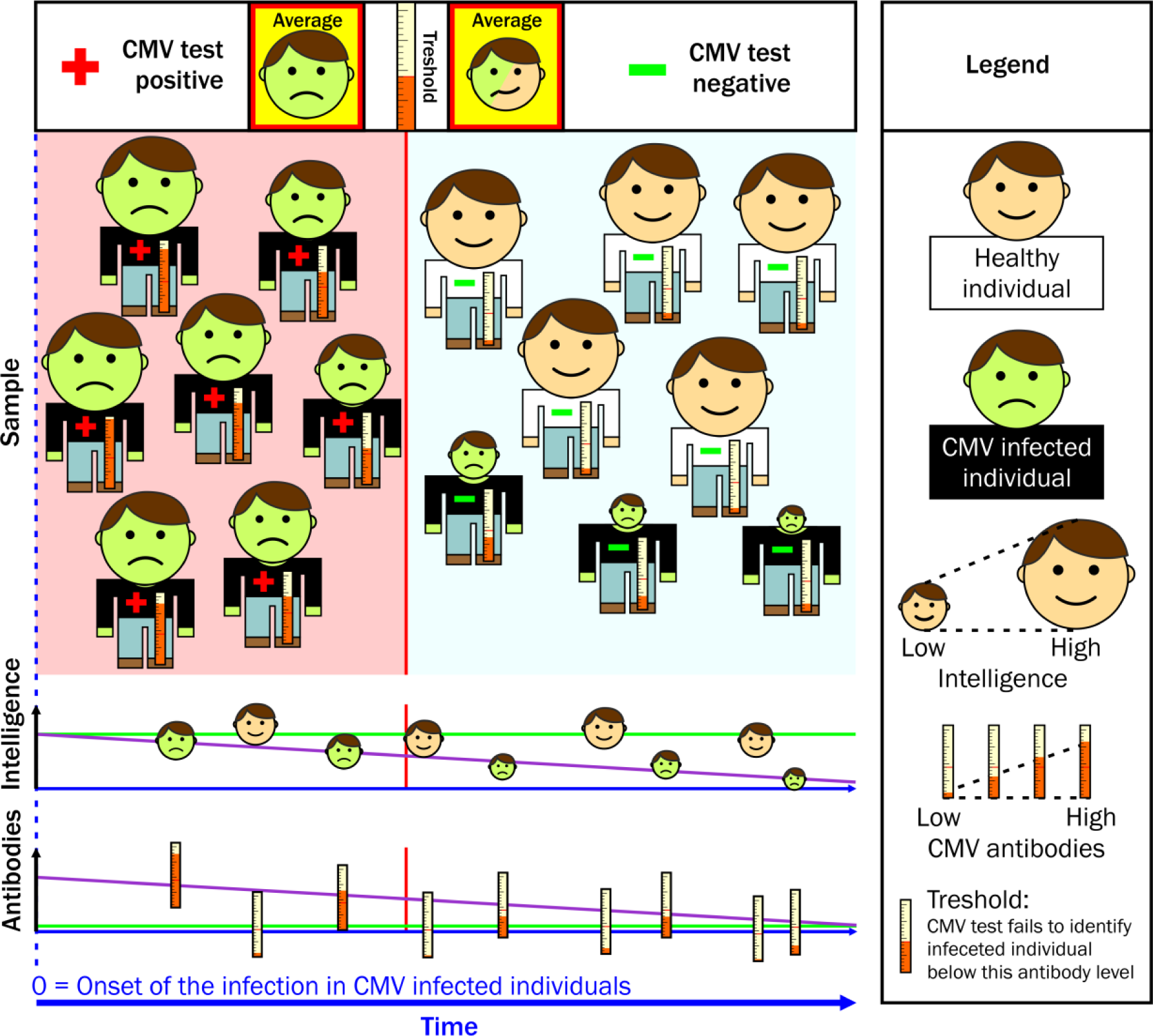
The model explaining the paradox of seemingly higher intelligence in CMV infected subjects and positive correlation between the concentration of anti-CMV antibodies and IQ.

The concentration of anti-CMV antibodies fluctuates in time and differs between subjects depending on many genetic and environmental factors, however, statistically, the level of these antibodies decreases with time from the infection in young people (see the lowest graph). In parallel, the intelligence of infected subjects decreases with time due to unknown cumulative effects of the chronical infection. Due to these two processes, the infected subjects with the lowest intelligence have also the lowest level of antibodies and many of them therefore score negatively in ELISA test (upper part of the figure). Therefore, the mean intelligence of CMV-seropositive subjects is higher than that of CMV-seronegative subjects (who represent the mixture on CMV-free and CMV-very-long-infected subjects).

The results of the permutation tests with reassignment 0, 5, 10, 15, 20, and 25 percent of CMV-seronegative subjects with the lowest intelligence from the CMV-free to CMV-infected set show Tab. 4 (all subjects), Tab. 5 (women), and Tab. 6 (men). The results suggest that seemingly higher intelligence of CMV infected subjects could be explained by the presence as low as 5 % of false negative subjects in our CMV-free set. When the effect of false negative subjects was controlled, the CMV-infected women expressed lower verbal knowledge while the CMV-infected men expressed lower verbal intelligence, verbal knowledge, general knowledge, and crystallized intelligence than their CMV-free peers.

**Table 4.**
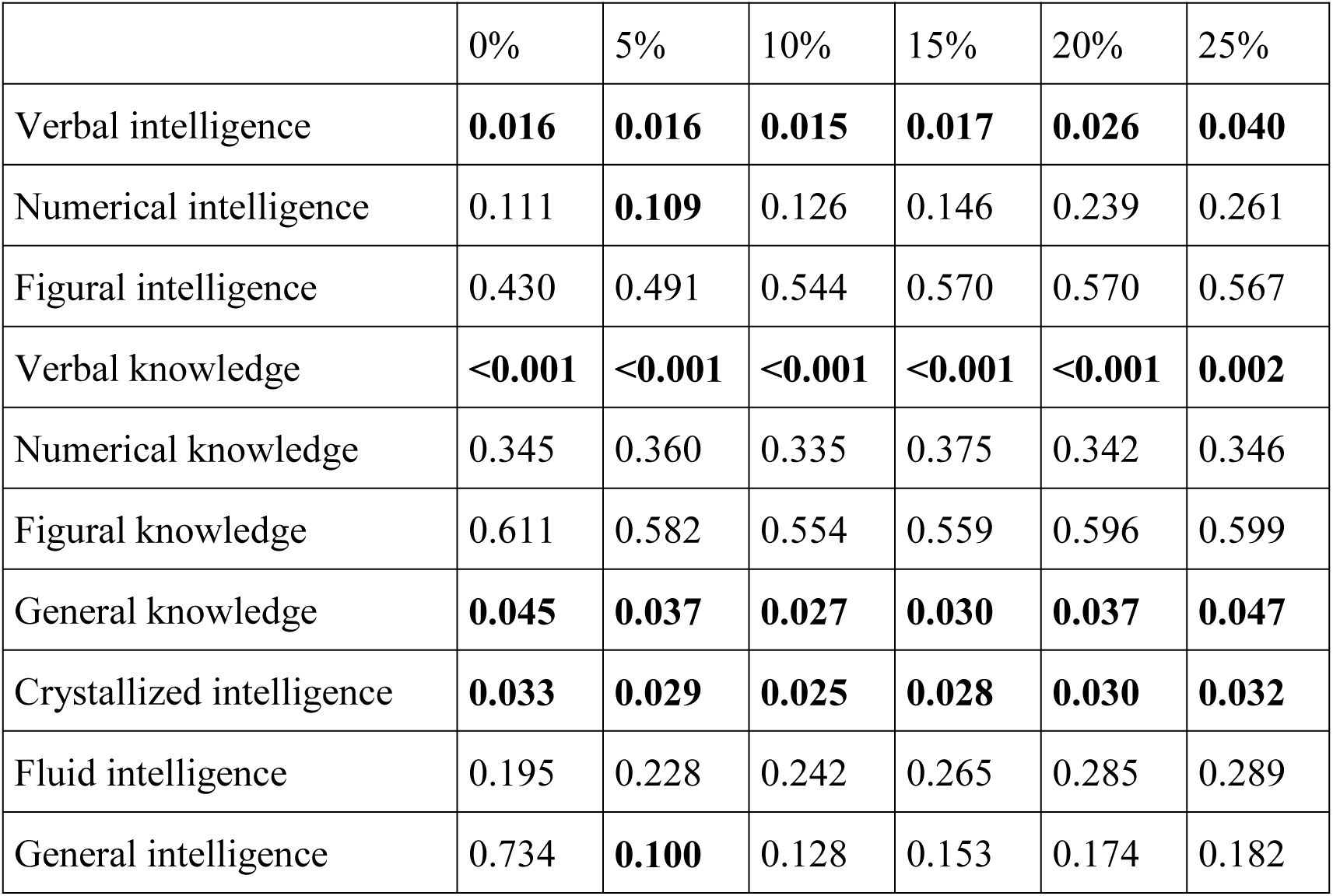
Differences between CMV-positive and CMV-negative subjects (women and men together) tested by permutation tail probability test with data reassignment.

The column headings show the fraction of reassigned cases and the rest of the table shows the statistical significance (p) of two-sided permutation tests. The p in the second column (without reassignment) correspond to H_0_ hypothesis, Infected subjects have lower or equal intelligence than CMV-free subjects. “while the p values in other columns correspond to H_0_ hypothesis: “Infected subjects have higher or equal intelligence than CMV-free subjects.”
This means that the significant results presented in the second column indicated higher cognitive performance of CMV seropositive subjects (when the effect of false negatives was not controlled), while the significant results presented in other columns indicated higher cognitive performance of CMV seronegative subjects (when the effect of false negatives was properly controlled). Results significant after the correction for multiple tests are printed in bold. The algorithm of the permutation test with data reassignment is described in the Methods section.

**Table 5.** Differences between CMV-positive and CMV-negative women tested by permutation tail probability test with data reassignment.

|  | 0% | 5% | 10% | 15% | 20% | 25% |
| --- | --- | --- | --- | --- | --- | --- |
| Verbal intelligence | 0.917 | 0.082 | 0.089 | 0.112 | 0.296 | 0.339 |
| Numerical intelligence | 0.868 | 0.124 | 0.156 | 0.173 | 0.192 | 0.202 |
| Figural intelligence | 0.622 | 0.367 | 0.437 | 0.474 | 0.489 | 0.520 |
| Verbal knowledge | 0.998 | **0.003** | 0.054 | 0.041 | 0.046 | 0.046 |
| Numerical knowledge | 0.126 | 0.691 | 0.669 | 0.646 | 0.659 | 0.633 |
| Figural knowledge | 0.146 | 0.708 | 0.738 | 0.845 | 0.845 | 0.847 |
| General knowledge | 0.739 | 0.215 | 0.215 | 0.235 | 0.274 | 0.358 |
| Crystallized intelligence | 0.767 | 0.091 | 0.116 | 0.128 | 0.151 | 0.206 |
| Fluid intelligence | 0.802 | 0.228 | 0.238 | 0.261 | 0.261 | 0.254 |
| General intelligence | 0.911 | 0.128 | 0.154 | 0.167 | 0.178 | 0.193 |

*For the legend see the Table 4.*

**Table 6.** Differences between CMV-positive and CMV-negative men tested by permutation tail probability test with data reassignment.

|  | 0% | 5% | 10% | 15% | 20% | 25% |
| --- | --- | --- | --- | --- | --- | --- |
| Verbal intelligence | **0.065** | **0.061** | **0.054** | **0.048** | **0.043** | **0.037** |
| Numerical intelligence | 0.444 | 0.446 | 0.452 | 0.453 | 0.465 | 0.489 |
| Figural intelligence | 0.676 | 0.691 | 0.704 | 0.701 | 0.698 | 0.674 |
| Verbal knowledge | **0.003** | **0.003** | **0.003** | **0.001** | **0.001** | **0.002** |
| Numerical knowledge | 0.178 | 0.174 | 0.167 | 0.154 | 0.174 | 0.171 |
| Figural knowledge | 0.606 | 0.602 | 0.550 | 0.452 | 0.404 | 0.232 |
| General knowledge | **0.070** | **0.058** | **0.047** | **0.039** | **0.030** | **0.028** |
| Crystallized intelligence | **0.051** | **0.038** | **0.037** | **0.035** | **0.040** | **0.041** |
| Fluid intelligence | 0.477 | 0.473 | 0.485 | 0.491 | 0.488 | 0.506 |
| General intelligence | 0.369 | 0.364 | 0.362 | 0.378 | 0.402 | 0.433 |

*For the legend see the Table 4.*

### f) Performance of the permutition test on non-contaminated data

To show that our permutation test for contaminated data cannot provide false positive results, i.e., cannot provide a more significant result than a standard permutation test if no false negative subjects exist in the population under study, we performed a Monte Carlo simulation using R 3.3.1 for Windows. We generated a population of 150 CMV-free and 150 infected subjects (mean intelligence was 101.5 in the CMV-free group and 98.5 in the infected group – the between-group difference was 3, the population mean intelligence was 100). Subjects were normally distributed around group means with equal standard deviations (SD). We used different SDs (3, 6, 9, 12, 15, 30) corresponding to different effect sizes expressed by Cohen’s d (1, 0.5, 0.33, 0.25, 0.2, 0.1). Then we ran a standard permutation test. We randomly permutated the CMV infection status of all subjects 10,000 times and computed in what fraction of permutations was the difference between two groups (pseudo-CMV-free and pseudo-CMV-infected subjects) equal or larger than the difference between the non-permutated data (p value of a standard permutation test). Then, we repeated the analysis using the permutation test for contaminated data. Namely, after generation of sets of CMV-free and CMV-infected subjects (or after generation of sets of pseudo-CMV-free and pseudo-CMV-infected subjects by permutation of CMV infection status), we relocated 5, 10, 15, 25 or 50% of subjects with the lowest intelligence from the CMV-free (or pseudo-CMV-free) set to the CMV-infected (or pseudo-CMV-infected) set. Again, we computed in what fraction of permutations was the difference between two groups equal or larger than the value computed for the non-permuted data (p values of the permutation test for contaminated data). We used populations generated for the standard permutation test (each initial population was used once for each fraction of relocated subjects). In total, 10,000 original populations were generated for each SD, therefore 10,000 independent permutation tests were conducted for each combination of SD and each relocated fraction. Resulting p values were averaged over permutation tests with the same population SD and relocated fraction and are reported in Table 7.

**Table 7.**
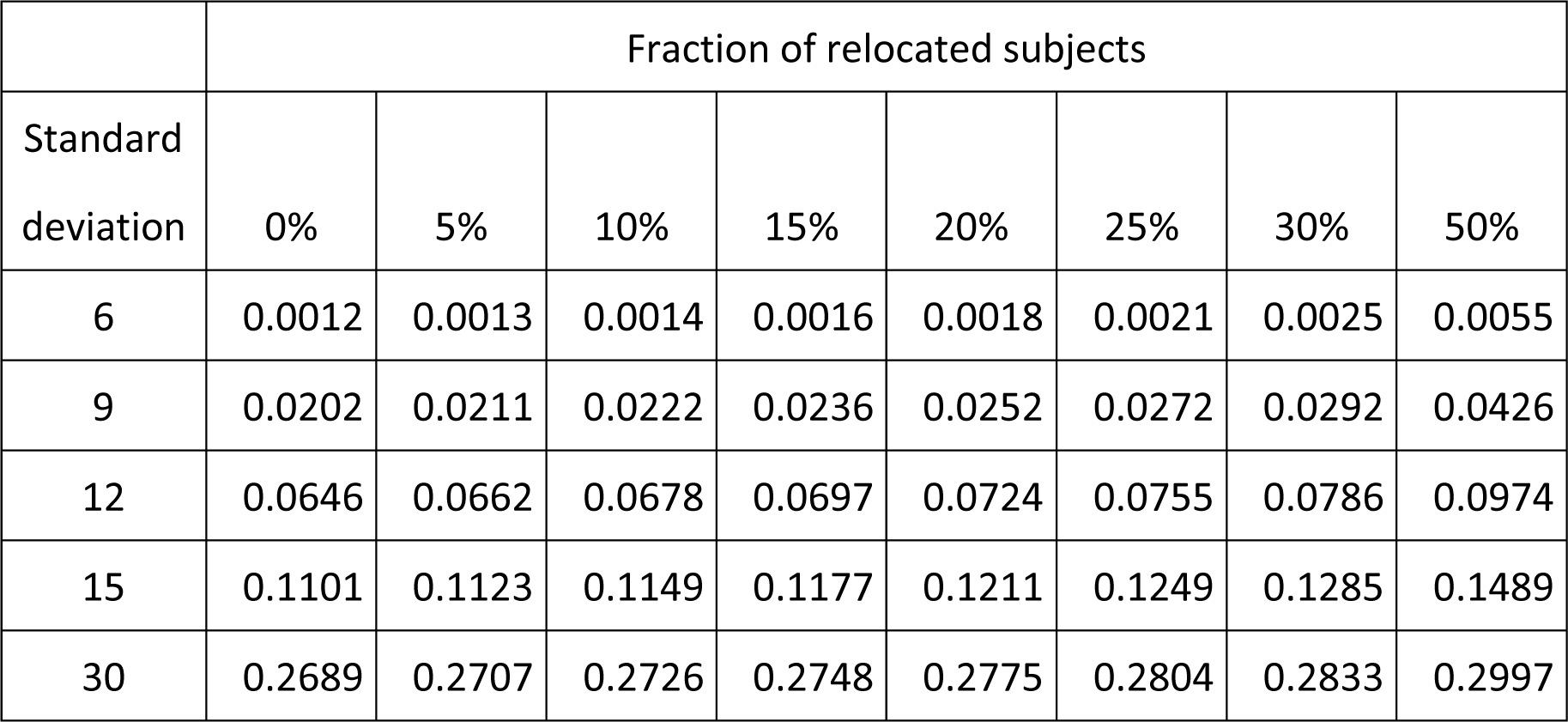
Effect of relocation of false negative subjects on the results of a permutation test if no such subjects exist in the population.

The table shows p values computed with the permutation test for contaminated data when the population under study contains no false negative subjects. The simulation experiments were performed on populations that differ by variances (rows) with relocation of different fractions of IQ-lowest individuals (columns) from the high-IQ group to the low-IQ group. The first column (0%) shows (the most significant) results of permutation tests performed without any relocation of data.

The Table 7 shows that the p value obtained with a standard permutation test is always lower than the p value obtained with the permutation test for the contaminated data when no subpopulation of false negative subjects exists in the population under study.

## Discussion

Students with anamnestic anti-CMV antibodies expressed higher intelligence, especially higher verbal knowledge. In the same time, the intelligence of CMV-positive subjects declined with the decrease of concentration of specific antibodies, which can be used (statistically, not in a clinical praxis) as a proxy of the time since the infection. The results of permutation test suggest that this paradox can be caused by the presence of as low as 5 % false negative subjects in the CMV-seronegative subset. Such false negative subjects in our population of students had probably the oldest infections and therefore also the lowest intelligence. Due to their presence in the seronegative subset, the mean intelligence of seronegative students was lower than the mean intelligence of seropositive students.

Our results suggest that CMV affects fluid, crystallized, and verbal intelligence and verbal knowledge. Fluid intelligence is a capacity to think logically and solve problems in novel situations, independent of cultural influences, dependent on biological cognitive prerequisites ^33^. It peaks in young adulthood and then steadily declines. Crystallized intelligence is the ability to use skills, knowledge, and experience; it is influenced by culture and education and it increases gradually with age ^33^. Verbal knowledge was the only component of intelligence affected in the subsample of women; however, verbal knowledge, verbal intelligence, crystallized intelligence, and general knowledge were affected in men and in the total sample. Therefore, the influence of CMV on intelligence of our subjects, especially the male subjects, is relatively nonspecific. Possibly, the worse performance of CMV-infected subjects in the intelligence tests can be caused by the same general defects, such as attention, learning and recall, which were already observed in healthy CMV seropositive middle-age adults ^16^.

We cannot even exclude possible effects of CMV infection on the motivation of subjects or their cooperativeness. It was, for example, already reported that CMV-infected subjects have lower novelty seeking and changed harm avoidance measured with Cloninger Temperament and Character Inventory ^17^. It is important to measure these variables in future studies to distinguish whether the intelligence or just the performance in the IQ test differs between CMV-infected and CMV-free subjects.

The existence of statistical association between two factors, here CMV infection and changed intelligence, can be explained in two principally different ways. Either the subjects with lower and higher intelligence differ in the probability of acquiring CMV infection or CMV infection influences the intelligence of infected subjects. The existence of positive correlation between specific anti-CMV antibodies and intelligence of CMV-positive subjects when the age of the subject is statistically controlled, however, suggests that the latter explanation, i.e., slow cumulative effect of CMV infection on human intelligence, is more parsimonious. It should be noted, however, that no cross sectional study can exclude the third alternative, namely, that less intelligent subjects acquire CMV infection earlier than subjects more intelligent.

The observed association of CMV with intelligence of seropositive subjects could be mediated by a third factor related both to CMV status and cognition. CMV seropositivity and higher CMV IgG titres in CMV-positive individuals were associated with lower leukocyte telomerase activity ^34^ and shorter leukocyte telomere length ^35^ in healthy adults. Longer telomeres, in turn, were shown to be associated with better cognitive performance in a large meta-analysis ^36^.

The present data agree with results of a previous study performed on military personnel^17^. This study, performed on 533 conscripts, demonstrated the higher intelligence (IQ = 97.8 vs 96.3, nonsignificant) of CMV-infected subjects and the positive correlation between concentration of anti-CMV antibodies and intensity of changes in personality in the infected conscripts. In that study, only the Otis test of verbal intelligence (a standard verbal intelligence test that consists of 32 questions focused on understanding of the given relationships, linguistic sensitivity, and vocabulary skills) was used and the age of subjects was not controlled.

Our results are also in agreement with results obtained with another pathogen, the *Toxoplasma gondii*. The life cycles of *Toxoplasma* and CMV differ, however, postnatal acquired infection with both *Toxoplasma* and CMV results in latent but most probably life-long infection of certain subpopulations of cells (probably including glial cells and monocytes in the case of CMV) that were considered harmless or even asymptomatic for a long time. Many studies published in past 20 years, nevertheless, showed that toxoplasmosis induces specific changes in behavior of infected animals ^37-39^ and in behavior and personality of infected humans ^40^. The personality changes observed in *Toxoplasma*-infected humans were searched for and observed also in the CMV-infected humans. It was, for example, observed that both *Toxoplasma* -and CMV-infected subjects have lower Cloninger’s novelty seeking. In *Toxoplasma*-infected subjects, the decrease of this factor is most probably caused by increased concentration of dopamine ^41-44^. Both *Toxoplasma* ^26,45,46^ and CMV ^17^ were suggested to influence also intelligence of infected subjects. Here, however, the results are somehow controversial. Latent toxoplasmosis was usually associated with decreased intelligence in men and sometimes with increased intelligence in women. Probably, this can be explained by existence of false negative subjects with the most decreased intelligence and the most decreased level of anamnestic antibodies in the studied female population. Even in our small set of 24 subjects who had been tested two or more times, we found one individual who was seropositive in the first, but seronegative in the last test. Results of previous studies that used the permutation test for contaminated data, observed decrease of seroprevalence in male population after the age of 36 ^47^ as well as observations of a conversion of seropositive to seronegative individuals ^48^, indicate that such subpopulation always exist in large experimental samples and can qualitatively influence results of observational studies ^26,32^.

Previous studies investigating the relationship of CMV infection and cognitive functions were done predominantly on population of children, elderly adults, schizophrenics and HIV patients. The main body of literature concerns congenitally acquired CMV infection in children with symptoms after the delivery. These symptoms include microcephaly, lethargy, seizures, paralysis, chorioretinitis and hearing loss ^49^. Forty to 58 % of symptomatic children have permanent sequelae, such as cognitive deficit or mental retardation ^50^. However, even 6.5 % of children who are asymptomatic at birth can develop some type of cognitive or neurological impairment ^50^, as both case-control studies ^51,52^ and one longitudinal study ^53^ showed. The incidence of congenital CMV infection is very low in the Czech Republic, being estimated lower than 1 %. The incidence of subjects with congenital CMV infection is probably even lower in the university students than in general population. Therefore, maximal expected occurrence of 2 – 3 such individuals among our probands cannot be responsible for the observed statistical associations.

No association between CMV and cognition was usually observed in asymptomatically infected subjects ^54-57^. Temple et al. ^58^ observed differences between infected and controls only in the group of younger children but not in the group of older children. Early postnatal infected subjects performed worse than controls both in very preterm infants ^59^ and in term infants ^60^. Even if there would not be any association between cognition and CMV in asymptomatically infected children as some authors indicated, it is possible that the subjects develop sequel later in life as research done on elderly adults suggests. Individuals with higher levels of IgG anti-CMV antibodies experienced more rapid decline of cognition over 4 year compared to subjects with lower levels of antibodies in a large group of CMV-infected elderly adults ^11^. In this case, higher levels of antibodies refer more probably to more frequent reactivations of the infection over the life course ^4^ rather than to a recently acquired infection. Similarly, in the seropositive subsample of older adults, higher level of antibodies to CMV was associated with lower general cognitive ability and processing speed. Moreover, CMV-positive subjects had lower cognitive ability than CMV-free controls ^10^. Another studies showed that cognitive functioning decreases with increasing viral burden including cytomegalovirus ^9,61^.

Our results are in agreement with those obtained on populations with pre-existing clinical conditions, on schizophrenia and on AIDS patients. CMV-infected individuals scored worse than those CMV-free in Trail Making Test in a set of schizophrenic patients^12^. Similarly, in a combined group of schizophrenics and controls the CMV – infected subjects performed worse than those uninfected as measured by the Wisconsin Card Sorting Test ^13^. Goplen et al. ^14^ conclude that CMV can act as a cause or an important cofactor of dementia symptoms in AIDS patients. Moreover, Lin et al. ^62^ found much higher percentage of CMV DNA in brains of individuals with vascular dementia compared to controls in an elderly sample.

Watson et al. ^15^ compared performance of patients with schizophrenia / schizo-affective disorder, their unaffected relatives and healthy controls. The authors observed worse performance of CMV-infected subjects compared to uninfected controls in all three groups, moreover, the differences were more pronounced in individuals with multiple infections (herpes simplex virus [HSV] 1, herpes HSV-2, CMV). Cognitive impairment was also reported for healthy middle-age either CMV or HSV-1 seropositive adults ^16^.

It must be reminded that no correction for presence of false negative subjects in the sample was performed in previous studies. Our results indicate that such subjects are probably often present in study populations and that as low as 5 % of such subjects can obscure the effect of CMV on intelligence. Therefore, proper measures against their effect on results of tests (such as use of permutation tests for contaminated data) should be used in future case-controls studies.

The number of experimental subjects (283) was relatively large; however, because of highly imbalanced sex ratio in the Charles University biology students, the number of men was only 86. This increased risk of Type II errors and also decreased the efficiency of controlling for potential confounding variables like the size of settlements where subjects spent their childhood, BMI, smoking, family background, or other (e.g. HSV-1, EBV, VZV, *Toxoplasma, Borrelia, Chlamidia, Candida*) infections. Most importantly, the effect of Rhesus D (RhD) phenotype that is known to strongly influence the effect of latent toxoplasmosis on human performance and personality ^46,63,64^ and also the effect of other factors (age and smoking) ^65,66^ on intelligence was not controlled in the present study. The Czech population contains only about 16 % RhD negative subjects. A much larger sample is thus necessary for searching for an RhD phenotype-CMV interaction or for controlling a broader spectrum of potential confounders to avoid the problem of over-parametrization of the models and of increased risk of false-positive or false-negative results of corresponding statistical tests..

In the present study, we used the concentration of anti-CMV IgG antibodies as a proxy of the time passed since the original CMV infection. This approach has already been used in other studies on different pathogens and is widely used in a clinical praxis, e.g. as an auxiliary screening technique for the prevention of congenital toxoplasmosis. It must be emphasized that the antibody level depends (possibly more strongly) also on many other factors (the infection dose, the virulence and antigenicity of the strain of a pathogen, number of successful or unsuccessful reinfections, the genetically or environmentally determined susceptibility of the host to the infection or to the resulting disease, etc.). Therefore, the relation between concentration of antibodies and time passed since the infection holds only statistically and probably only in certain phases of the infection. It must be emphasized that all previously mentioned factors, except the time passed since the infection, would result in the existence of a POSITIVE correlation between the level of antibodies and the observed effects of the infection. In the present study, however, we detected a NEGATIVE correlation between the antibody level and impairment of cognitive performance of the students. This suggests, but of course does not prove, that the antibody level in the biology students most likely (statistically) reflects the time passed since the infection, and not, for example, the intensity or frequency of past infections.

Last but not least, the biology students of the most prestigious Czech university, Charles University in Prague, are not typical representatives of a general Czech population. Therefore, it is not clear to what extent the observed phenomena can be generalized. We could expect to detect a much stronger effect of CMV on an “unsorted” population that had not passed recently through a sieve of relatively severe entrance examinations. This sieve probably eliminated a larger fraction of low-IQ (and therefore the CMV seropositives-enriched) subjects than that of high-IQ (the CMV seropositives-impowerished) subjects. Among the CMV seropositives, only the individuals who had a very high IQ before the infection could successfully pass through the examinations while among the CMV seronegatives, also the less gifted subjects could pass the same examinations. If the entrance examinations work properly, the IQ of CMV seropositive and seronegative students would be very similar immediately after the entrance examination but the representation of CMV seropositive individuals would be lower in successful examinees (which could be tested, of course). It will therefore be more effective to study the effect of CMV seropositivity (or of any other environmental factors) on the representatives of a general population or at least on subjects before, not after, the entrance examinations or any similar “sieves”.

Also, the existence of the strict sieve effect of the entrance examinations could explain observed difference in intelligence between male and female students. The mean intelligence of men and women in general population is approximately the same, however, the spread of the intelligence is larger in men than in women. In contrast to the situation in the general population, higher mean intelligence in women than men can be therefore expected in any subpopulation of low intelligence subjects (e.g. students of less prestigious schools) and higher mean intelligence in men than women can be expected in any subpopulation of high intelligence subjects (e.g. students of the most prestigious universities).

An unknown third factor, e.g. socioeconomic situation of a subject, could correlate both with probability of CMV infection and the intelligence. Czech population and especially the university students have extremely low socioeconomic stratification. Still, broader spectrum of potential confounding variables should be monitored and controlled for in future studies.

## Conclusions

The present study performed on nearly three hundred university students showed that CMV infection is associated with seemingly increased intelligence measured with The Intelligence Structure Test I-S-T 2000 R. The negative correlation between intelligence and duration of the infection estimated on the basis of concentration of anti-CMV antibodies and the results of the permutation test suggests that the seemingly increased intelligence of CMV-infected subjects is a result of presence of the subpopulation of subjects with very old infection and therefore also the most decreased intelligence in the subpopulation of seronegative subjects. In accordance with the Bradford-Hill 5^th^ criterion of causality, this negative correlation between intelligence and concentration of anti-CMV antibodies also suggests, but, of course, not definitively proves, that the CMV-intelligence association is rather the effect than the cause of the CMV infection. CMV infection in both developing and developed countries is very high; most of the world population is probably infected with this herpetic virus. Therefore, the total impact of CMV on human intelligence, and secondarily on all the other aspects of human life, including quality of life and economy, may be extremely high.

## Acknowledgements

We would like to thank Lasha Lanchava, Zdeněk Hodný and Lenka Priplatova for their advices on the draft of this paper. The work was supported by project UNCE 204004 (Charles University in Prague) and the Czech Science Foundation (Grant No. P303/16/20958).

## Authors’ contributions

JF designed the study and wrote the article, VC performed research, analyzed the data and wrote the article, BS, PT, and HH performed research and participated in writing the article.

